# Loss of Tonoplast Sucrose Transporter SUT4 Alters Seasonal Growth, Carbohydrate Allocation, and Fertility in Field-Grown Poplar

**DOI:** 10.1101/2024.10.22.619524

**Authors:** Trevor T. Tuma, Holly A. McInnes, HongDuyen Pham, William P. Bewg, Michihito Deguchi, Ran Zhou, Samantha M. Surber, Brent Lieb, Anna Lipzen, Kerrie W. Barry, Daniela L. Weber Wyneken, Anne E. Harman-Ware, Joseph Dahlen, Scott A. Harding, Chung-Jui Tsai

## Abstract

Climate uncertainty is intensifying the need for greater plasticity in carbohydrate reserve utilization to support winter survival and spring growth in woody perennials. In poplar, the single-copy *SUT4*, which encodes a tonoplast-localized sucrose transporter, and the *SUT5*/*SUT6* genome duplicates, which encode plasma membrane-localized transporters, are expressed year-round, with *SUT4* showing highest expression during cool seasons. Given its role in vacuolar sucrose efflux and winter-predominant expression, SUT4 may play a key role in modulating seasonal carbohydrate dynamics. While *SUT4-*knockdown and knockout effects have been studied under greenhouse conditions, their impact under field conditions remains unexplored. Here, we report a field-based study comparing CRISPR knockout mutants of winter-expressed *SUT4* and *SUT5*/*SUT6* in *Populus tremula* × *alba*. We show that *sut4*, but not *sut56*, mutants exhibited earlier autumn leaf senescence, delayed spring bud flush, reduced stem growth, and altered sugar partitioning in winter xylem and bark relative to controls. After two years in the field, all genotypes flowered before leaf flush in early spring; however, *sut4* mutants produced sterile ovules despite developing normal-looking catkins. Metabolic profiling revealed disrupted sucrose and raffinose dynamics in elongating *sut4* catkins, accompanied by transcriptomic signatures of elevated stress and downregulation of proanthocyanidin biosynthesis and circadian clock genes. These findings highlight the critical role of SUT4 in coordinating sugar allocation, stress responses, and seasonal development in poplar.

**Significance statement:** This field study demonstrates that loss of SUT4, the most highly expressed sucrose transporter during cool seasons, disrupts phenology, growth, and fertility in poplar. Altered sugar and raffinose dynamics and transcriptomic signatures of stress and circadian clock gene dysregulation in the mutants underscore SUT4’s role in coordinating sugar allocation and seasonal developmental transitions under natural environmental conditions.

## Introduction

Woody perennials in temperate regions undergo an annual cycle of growth, dormancy, and resumption in response to seasonal cues such as temperature and day length. Following autumnal leaf drop, these species rely on stored carbohydrate reserves to support critical transitions through dormancy, floral development, flowering, and leaf emergence in the spring (Rohde and Bhalerao, 2007, Cook *et al*., 2012, Savage and Chuine, 2021). Rising temperatures and increasing climate variability are exacerbating uncertainty in the timing, trajectory, and phenological consequences of seasonal weather transitions (Tanino *et al*., 2010, Rohde *et al*., 2011, Calinger and Curtis, 2023). For perennial plants, this uncertainty underscores the growing importance of plasticity in carbohydrate resorption, storage, and remobilization for long-term resilience.

An important player in this context may be the tonoplast sucrose transporter SUT4, which mediates sucrose efflux from the vacuole into the cytosol (Reinders *et al*., 2008, Payyavula *et al*., 2011, Schneider *et al*., 2012). *SUT4* expression is sensitive to drought and temperature stress, and its regulation by hormone and light quality signals positions it as an integrative component in carbohydrate transport plasticity (Chincinska *et al*., 2013, Gong *et al*., 2015, Xue *et al*., 2016, Xu *et al*., 2017). In *Populus*, *SUT4* is expressed at much higher levels in overwintering than in summer stems and buds (Ko *et al*., 2011, Sreedasyam *et al*., 2023, Tuma *et al*., 2024), suggesting a potential seasonal role. Based on evidence of its importance for hydrostatic control and abiotic stress tolerance in various organs during active growth (Frost *et al*., 2012, Xue *et al*., 2016, Harding *et al*., 2020, Harding *et al*., 2022), SUT4 may also serve nutritive and/or protective roles in overwintering trees.

A poplar coppicing study using *SUT4*-knockout (KO) mutants provided evidence for a role of SUT4 in the emergence and early growth of epicormic buds under suboptimal winter glasshouse conditions (Tuma *et al*., 2024). Although starch and sucrose depletion magnitudes were similar between *sut4* and wild-type (WT) plants, epicormic bud release and sprout growth were reduced in the mutants, consistent with impaired vacuolar sucrose efflux (Tuma *et al*., 2024). Notably, this coincided with higher raffinose abundance in the winter stem of *sut4* and a steeper decline following coppicing relative to WT. Raffinose, a trisaccharide synthesized directly from sucrose in the cytosol (Schneider and Keller, 2009), is known to contribute to membrane stabilization, reactive oxygen species (ROS) scavenging, and protection against winter freezing (Ameglio *et al*., 2004, Regier *et al*., 2010). Raffinose accumulation is modulated by multiple abiotic stress signaling pathways (Pluskota *et al*., 2015, Khan *et al*., 2021, Noronha *et al*., 2022), which may explain why its abundance does not consistently align with *SUT4* expression in *SUT*-KO or knockdown (KD) poplars (Frost *et al*., 2012, Harding *et al*., 2022, Tuma *et al*., 2024). Furthermore, storage water capacitance and modulus of elasticity differ between WT and *SUT4*-KD lines (Harding *et al*., 2020). During drought stress, trajectories of leaf relative water content reductions also differ between WT and *SUT4*-KO or KD lines (Harding *et al*., 2022). These findings suggest that the metabolic outcomes of SUT4 plasticity are shaped by multiple stress signaling pathways—some that directly modulate *SUT4* expression (Gong *et al*., 2015), and others that are themselves modulated by SUT4 activity (Xue *et al*., 2016).

Under non-stress conditions, plant growth, cell wall composition, and vessel development were altered in transgenic poplar with elevated raffinose levels via overexpression of a *galactinol synthase* gene (Unda *et al*., 2017). Although not necessarily raffinose-dependent, spring bud break and growth resumption are partially influenced by stem vessel architecture, the timing of new xylem formation, and post-winter hydraulic capacity (Lechowicz, 1984, Perrin *et al*., 2017). More broadly, raffinose may modulate salicylic acid signaling (La Mantia *et al*., 2017), which in *Populus* has been shown to affect carbon allocation toward flavonoid-derived proanthocyanidins (*i.e.*, condensed tannins) with documented protective functions in leaves (Gourlay and Constabel, 2019, Ullah *et al*., 2019). Proanthocyanidins also play protective roles during reproductive development (Debeaujon *et al*., 2000, Dixon and Sarnala, 2020).

In poplar, vascular-specific Type I *SUT* transcripts (*SUT1* and *SUT3* genome duplicates) are detected only in the summer, whereas the broadly expressed Type II (*SUT5* and *SUT6* genome duplicates) and Type III (single-copy *SUT4*) transcripts are present year-round, with *SUT4* predominating during the cool seasons (Ko *et al*., 2011, Sreedasyam *et al*., 2023, Tuma *et al*., 2024). The present study investigated the effects of *SUT4*-KO under field conditions. For comparison, we also generated single- and double-KO mutants of the other winter-expressed *SUT5/SUT6* paralogs encoding plasma membrane transporters (Peng *et al*., 2014). While our results indicate that *SUT5/SUT6*-KO (hereafter *sut56*) had no observable effects on seasonal growth or carbohydrate metabolism, *SUT4*-KO (*sut4*) mutants exhibited accelerated leaf senescence in the fall and delayed bud flush in the spring, culminating in reduced biomass growth. The field study further revealed SUT4-dependent effects on the metabolic trajectories of developing female catkins. Together, these findings strengthen the rationale of SUT4 as a key modulator of sugar allocation and stress response mechanisms during seasonal growth transitions, with implications for perennial plant adaptation to changing climates.

## Results

### Altered phenology and reduced stem growth in sut4 but not sut56 mutants

We established a field trial in summer 2021 to investigate the effects of winter-expressed *SUT* gene knockouts on seasonal growth and phenology. Over 200 trees were planted in 20-gal pots; cohort 1 (control and *sut4*) in early June and cohort 2 (control, *sut4*, *sut5*, *sut6*, and *sut56*) in mid-August. In late October, following their first (partial) growth season, *sut4* mutants showed signs of leaf yellowing while leaves of the other genotypes remained green regardless of cohort (Fig. 1a). By mid-November, nearly 90% of *sut4* trees were in a denuded state, whereas the majority of the controls and *sut5*, *sut6*, and *sut56* mutants still retained most of their foliage (Fig. 1b-c). Post-dormancy bud flush the following spring (March 2022) was delayed by an average of 7 days in *sut4* trees compared to the control and *sut5/sut6/sut56* mutants (Fig. 1d). Similar trends were observed in two subsequent years, suggesting a reduced leaf lifespan in *sut4* trees.

**FIGURE 1:**
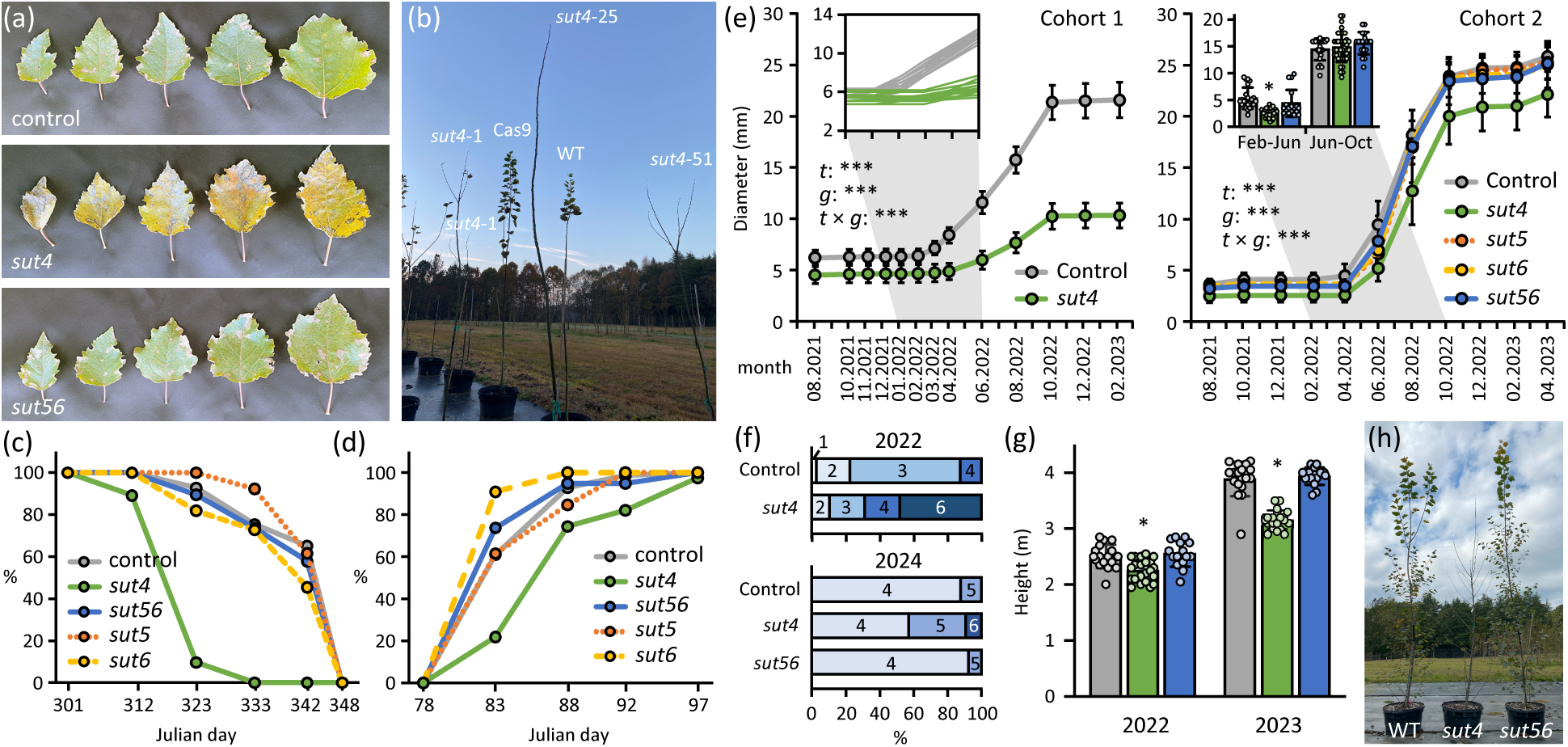
Biomass yield and growth characteristics of *sut* and control poplars. (a) Representative adaxial views of leaf plastochron index (LPI)-4, LPI-6, LPI-8, LPI-10, and LPI-12 prior to leaf abscission. (b) Representative photos of field-grown trees taken in November 2021. (c) Progression of leaf abscission from the main stem in Fall 2021. Data represent the percentage of trees per genotype that had abscised or desiccated leaves using pooled transgenic events and biological replicates (*n =* 69 for control, *n =* 78 for *sut4*, *n =* 13 for *sut5*, *n =* 11 for *sut6*, and *n =* 19 for *sut56*). (d) Bud flush was evaluated in the spring of 2022 based on the epicormic buds on the main stem. Data represent the percentage of trees per genotype that had flushed buds using pooled transgenic events and biological replicates (*n =* 69 for control, *n =* 78 for *sut4*, *n =* 13 for *sut5*, *n =* 11 for *sut6*, and *n =* 19 for *sut56*). (e) Stem diameter of trees from cohort 1 and cohort 2 measured 1 m above root collar monthly or bi-monthly. Data are mean ± standard deviation of pooled transgenic events and biological replicates (*n =* 40 for control or 29 for *sut4* in cohort 1, or *n =* 23 for control, 40 for *sut4*, 13 for *sut5*, 11 for *sut6*, and 18 for *sut56* in cohort 2). The effects of time (*t*), genotype (*g*), and their interaction (*t* × *g*) were assessed with a repeated measures two-way ANOVA. ***, *P* ≤ 0.001. Inset in cohort 1 was from size-matched control or *sut4* trees at the start of the monitoring (*n* = 13). Inset in cohort 2 represent diameter increments during spring and summer of 2022. (f) Cambial growth reactivation in the spring of 2022 (cohort 1) and 2024 (cohort 2). Data represent the percentage of trees per genotype that had the first measurable diameter increases, with months indicated by numbers (*n* = 29-40 trees for cohort 1 or *n* = 13-24 trees for cohort 2). (g) Height of dominant stem measured at the beginning of the growing season in 2022 and 2023. Data are mean ± standard deviation of *n =* 15 – 22 trees per genotype using pooled transgenic events and biological replicates. Significance was conducted using 2-sample *t-*test. *, *P <*0.05. (h) Representative image of control, *sut4,* and *sut56* trees 14 months after field outplanting.

Main stem diameter growth was monitored monthly for cohort 1 and bimonthly for cohort 2. Since no growth differences were observed among independent events across all traits, data were pooled by mutant for ease of comparison. Diameter growth in *sut5*, *sut6*, and *sut56* trees was indistinguishable from controls, whereas *sut4* trees exhibited significantly reduced growth during the 2022 season (Fig. 1e). This reduction was more pronounced in cohort 1 than in cohort 2, consistent with previous reports of poor growth performance following summer outplanting (Grossnickle, 2005, Landhäusser *et al*., 2012). In cohort 1, all control trees had resumed cambial growth and showed increased diameters by April 2022, whereas only 51% of *sut4* trees exhibited measurable growth (Fig. 1f). To determine whether the delayed spring growth in *sut4* trees was due to their smaller initial size, we analyzed 13 size-matched control and *sut4* trees. We found that *sut4* diameter growth still lagged by 2-3 month compared to size-matched controls (Fig. 1e cohort 1 inset). Similarly, in cohort 2, diameter growth of *sut4* trees between February and June 2022 was half that of control and *sut56* trees (Fig. 1e cohort 2 inset), through growth rates were comparable among genotypes from June to October 2022 (Fig. 1e cohort 2 inset). The 2022-2023 winter was mild (Fig. S1), and diameter differences between *sut4* and the other genotypes remained constant through the end of field trial in April 2023 (Fig. 1e). A subset of cohort 2 trees was permitted to continue, and by April 2024, diameter growth had resumed in 92% of control and *sut56* trees, but in only 58% of *sut4* trees (Fig. 1f).

Growth was unaltered in both the single- and double-KO of *SUT5/SUT6*; therefore, only *sut56* trees were included in subsequent analyses. Tree height was slightly reduced, by ∼10%, in *sut4* compared to the controls and *sut56* trees in April of 2022 (Fig. 1g-h). By April 2023, *sut4* trees were ∼ 20% shorter than control and *sut56* trees (Fig. 1g). Taken together, reduced growth, shorter leaf lifespans, and delayed spring cambial growth were consistently observed in *sut4* trees over multiple years.

### Altered nonstructural carbohydrate abundance in sut4 stems

We compared levels of monosaccharides and oligosaccharides in the bark and xylem tissues sampled during summer and winter seasons. Bark and xylem sucrose levels were higher in *sut4* mutants than the other lines during the summer (Fig. 2a), consistent with previous findings from greenhouse-grown plants (Payyavula *et al*., 2011, Harding *et al*., 2022, Tuma *et al*., 2024). Sucrose levels decreased during the winter most drastically in *sut4*, neutralizing most but not all of the summer genotypic increases (Fig. 2a). Fructose and glucose, the major stem hexoses, exhibited low abundances in all lines during the summer and increased in the winter by 1-2 orders of magnitude in both tissues, reaching highest abundances in *sut4* in both bark and xylem (Fig. 2b). Bark and xylem starch abundance differed little between genotypes and exhibited similarly large declines in all genotypes during winter (Fig. 2c).

**FIGURE 2:**
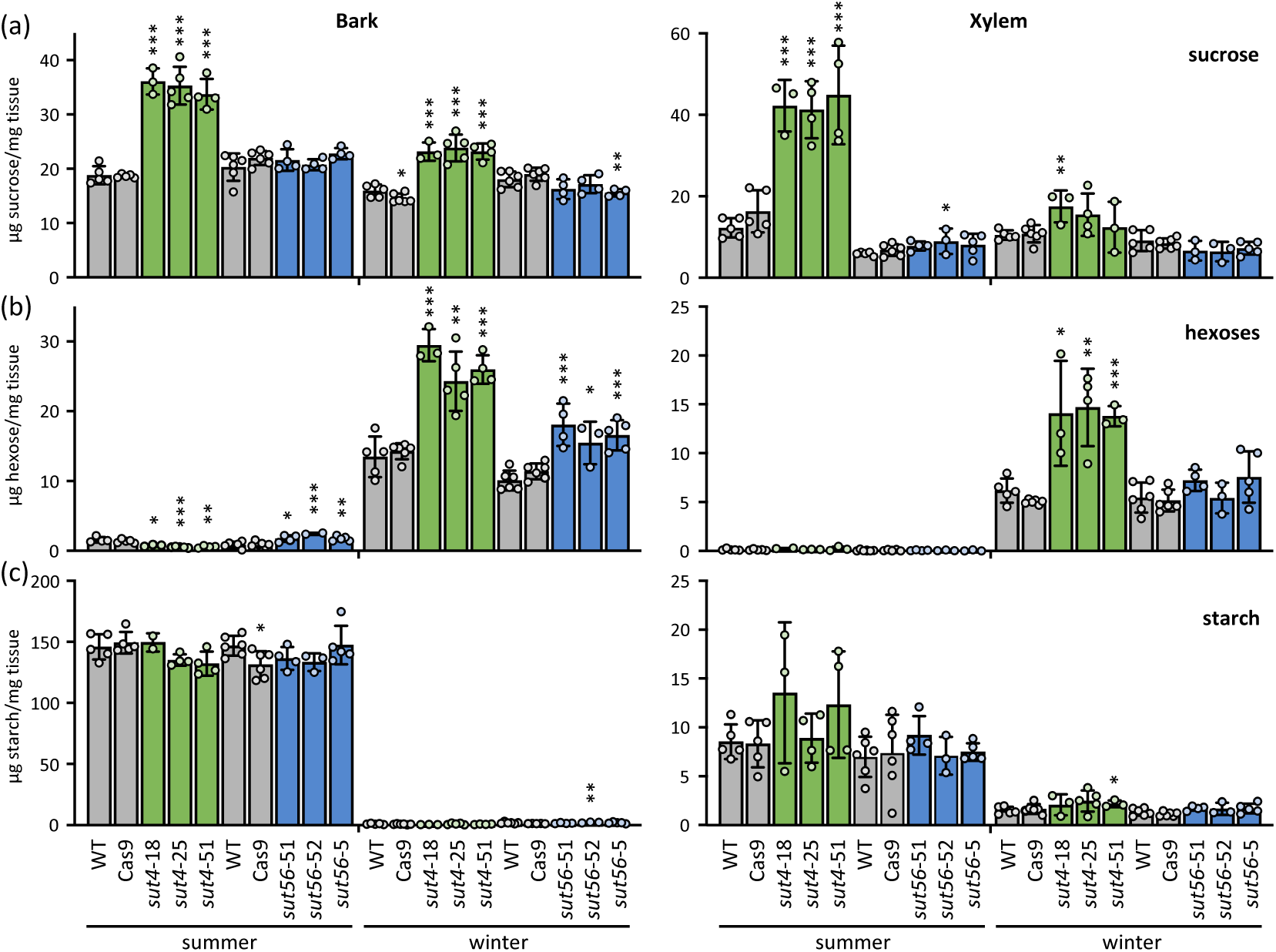
Nonstructural carbohydrate abundance in stem organs of different genotypes. (a) Sucrose, (b), hexose (sum of fructose and glucose), and (c) starch abundance in bark and xylem during summer and winter. Data are mean ± standard deviation of *n =* 3-6 biological replicates. Significance was determined by Student’s *t*-test against WT. ∗, *P* ≤ 0.05; ∗∗, *P* ≤ 0.01; ∗∗∗, *P* ≤ 0.001.

Raffinose and its precursor galactinol are known to accumulate under desiccation stress such as drought and cold (Yan *et al*., 2022). Galactinol and raffinose were only marginally detected in summer whereas their precursor myo-inositol was readily detected and significantly more abundant in both bark and xylem of *sut4* trees compared to other genotypes (Fig. 3a). During winter, galactinol and raffinose levels increased sharply in all genotypes, presumably at the expense of myo-inositol (Fig. 3a-c). However, these winter increases were significantly attenuated in *sut4* bark and trended so in xylem (Fig. 3b-c). Overall, stem sucrose and hexose data did not readily support carbohydrate provisioning *per se* as the cause of delayed spring bud flush or stem diameter growth in *sut4* trees. Nonetheless, alternations in carbohydrate metabolism, including raffinose dynamics, were observed under outdoor winter conditions and may be relevant to the growth phenotypes of *sut4* trees.

**FIGURE 3:**
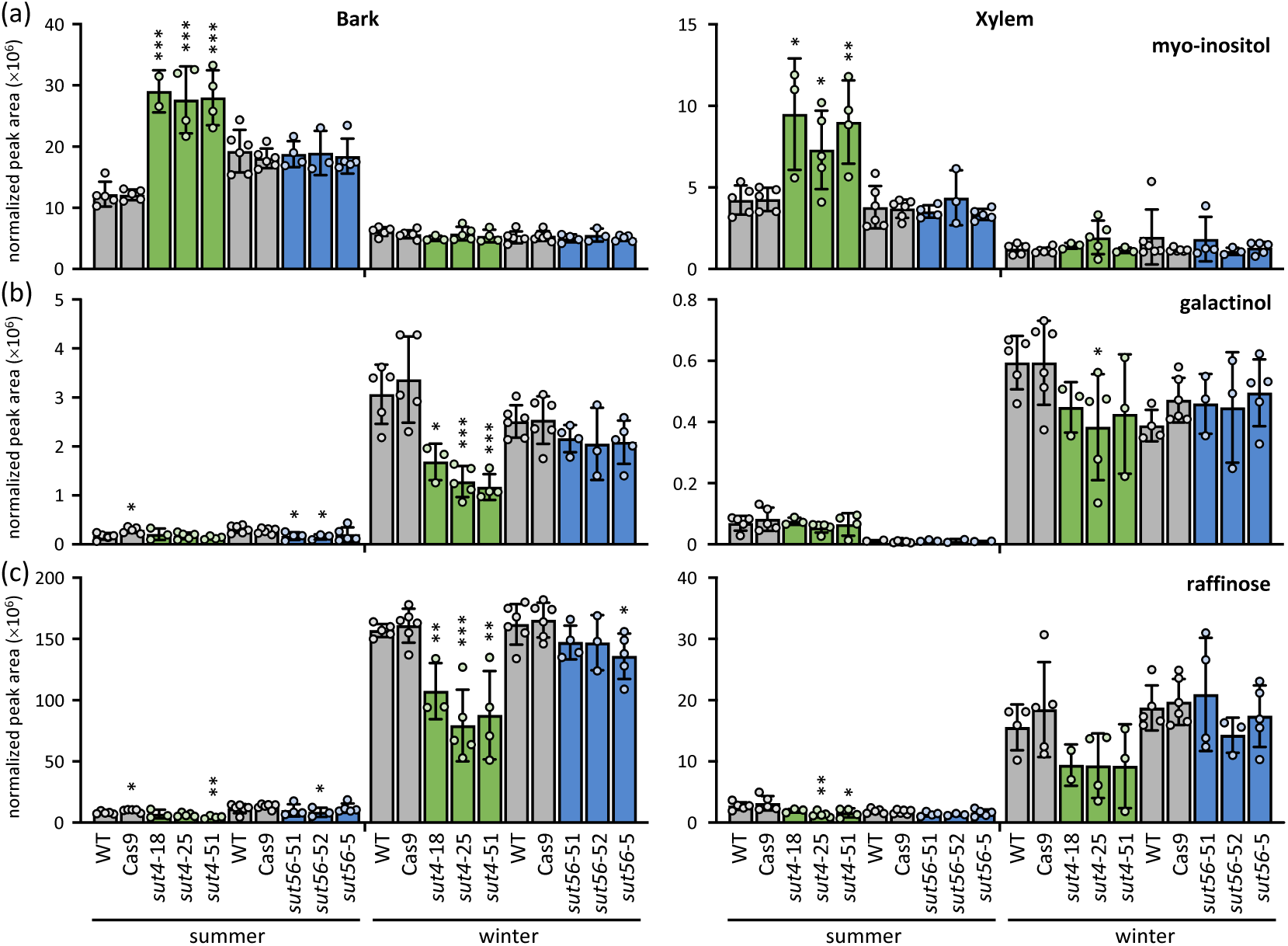
Oligosaccharide abundance dynamics in stem organs of different genotypes. (a) Myo-inositol, (b), galactinol, and (c) raffinose abundance in bark and xylem during summer and winter. Data are mean ± standard deviation of *n =* 3-6 biological replicates. Significance against WT was determined by Student’s *t*-test. ∗, *P* ≤ 0.05; ∗∗, *P* ≤ 0.01; ∗∗∗, *P* ≤ 0.001.

### Wood physicochemical properties were altered in sut4 trees

In light of the reduced stem diameter growth observed in *sut4* lines, wood samples were analyzed for changes in chemical and physical properties. Neither lignin content nor syringyl-to-guaiacyl (S/G) monolignol ratio was altered in *sut4* or *sut56* trees compared to controls (Fig. 4a-c). Acoustic velocity also did not differ between genotypes (Fig. 4d). However, specific gravity and the modulus of elasticity were reduced in *sut4*, but not *sut56*, relative to the controls (Fig. 4e-f). These findings suggest that the delay onset of spring diameter growth in *sut4* trees may reflect more complex effects of *SUT4*-KO on structural development either seasonally or year-round.

**FIGURE 4:**
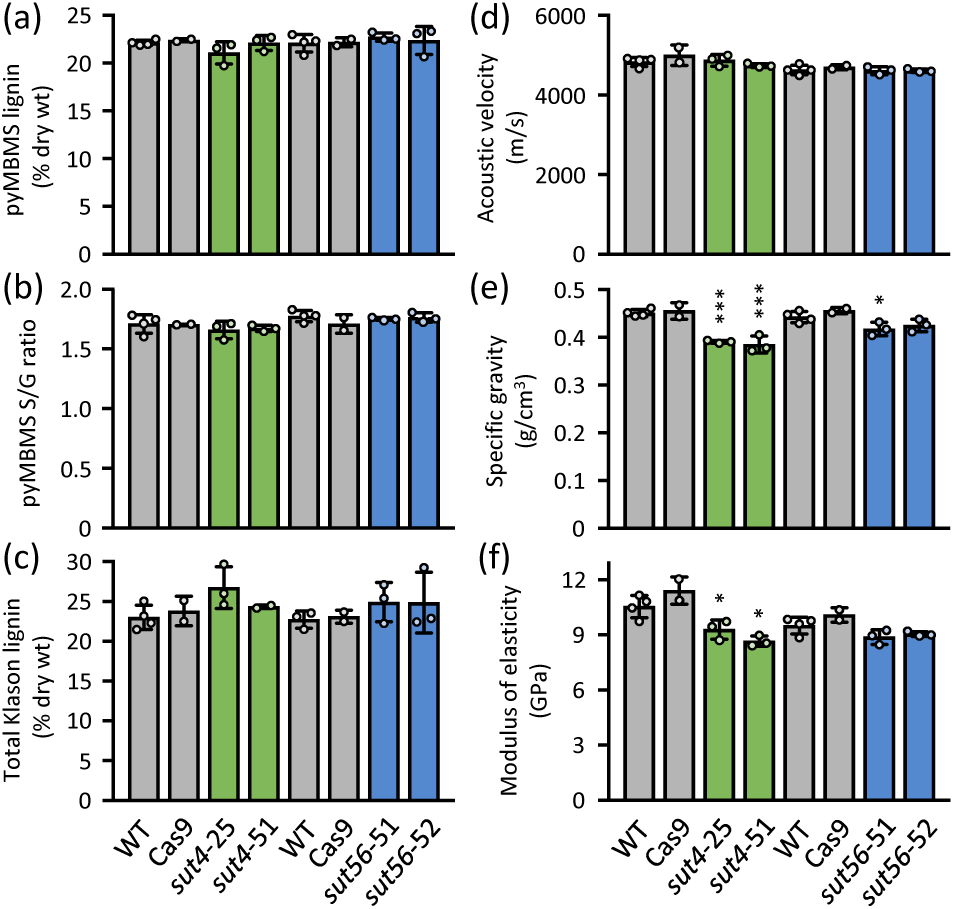
Wood characteristics and chemical properties. (a) Total lignin content and (b) syringyl-to-guaiacyl lignin (S/G) ratio determined by pyMBMS, (c) total Klason lignin content, (d) acoustic velocity, (e) specific gravity, and (f) modulus of elasticity. Data are mean ± standard deviation of *n =* 2-4 biological replicates. Significance was determined against WT trees by Student’s *t*-test; ∗∗, *P* ≤ 0.01; ∗∗∗, *P* ≤ 0.001.

### Reproductive growth and development were altered in sut4 trees

Field-grown female 717 trees typically flower after reaching 5-8 years of age (Mohamed *et al*., 2010). However, during the spring of year 2 and prior to leaf emergence, catkins were unexpectedly observed on many nursery trees (Fig. 5a). No consistent differences in catkin emergence frequency were observed between genotypes in either spring 2023 or 2024. In 2023, flowering frequency appeared to correlate with tree size: an average of 69% of cohort 1 trees produced catkins compared to 36% in cohort 2 which had been released two months later (Fig. 5b). In the extended trial with a subset of cohort 2 trees, 75% produced catkins in 2024, regardless of genotype (Fig. 5b).

**FIGURE 5:**
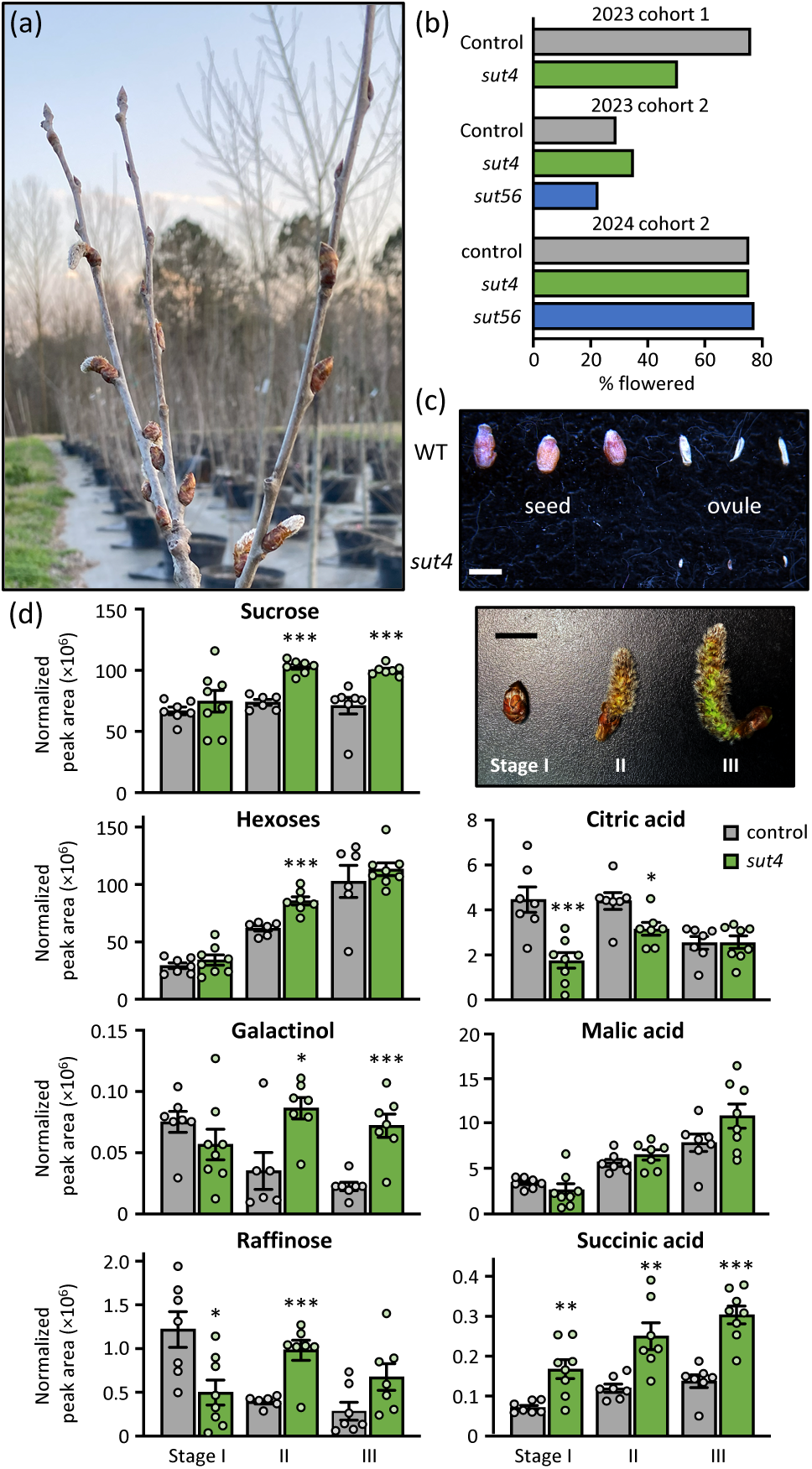
Flowering phenotypes in control and *sut* trees. (a) Axillary inflorescence bud flushing in early spring 2023. (b) Percentage of trees per genotype from Cohort 1 and Cohort 2 that developed catkins during spring of 2023 and 2024 (*n =* 41 control or 29 *sut4* trees for 2023 cohort 1; *n =* 27 control, 49 *sut4*, or 18 *sut56* trees for 2023 cohort 2; *n =* 24 control, 36 *sut4*, or 13 *sut56* trees for 2024). (c) Representative images of seeds and ovules from WT and *sut4* catkins. Scale bar = 1 mm. (d) Abundance of sucrose, hexose (sum of fructose + glucose), galactinol, raffinose, citric acid, malic acid, and succinic acid during catkin development. Representative developmental stages are shown in the photo (I, unopened floral buds; II, newly emerged catkins; and III, elongating catkins). Bar = 1 cm. Data are mean ± standard error of *n =* 6-8 biological replicates. Significance was determined by Student’s *t*-test. ∗, *P* ≤ 0.05; ∗∗∗, *P* ≤ 0.001.

To test whether *SUT4*-KO affected fertility, we performed controlled pollination using wild *P. tremuloides* pollen. Of the 13 WT and 14 *sut4* catkins pollinated, nine from each genotype developed capsules with abundant seed trichomes. While WT flowers produced seeds as expected, no seeds were recovered from *sut4* capsules (Fig. 5c). Although unfertilized ovules in aspens can undergo limited development (Fechner, 1972), the ovules in *sut4* capsules were notably smaller than those in WT (Fig. 5c). It is unclear whether the observed sterility in *sut4* trees was due to ovule failure, other developmental anomalies, or environmental sensitivity associated with SUT4 disruption.

### Metabolic profiling revealed contrasting developmental trajectories between WT and sut4 catkins

*SUT4* is the predominant *SUT* gene expressed in poplar flowers (JGI Plant Gene Atlas, Sreedasyam *et al*., 2023), suggesting that cytosolic access to vacuolar sucrose reserves is likely important during floral development. Given that the timing of ovule development relative to catkin growth is known (Wang *et al*., 2012), we reasoned that metabolic profiling could reveal how loss of SUT4 function impacted fertility. Metabolic profiling was conducted on unopened floral buds, newly emerged catkins, and elongating catkins, designated as stages I, II, and III, respectively (Fig. 5d). The temporal dynamics of sucrose abundance differed markedly between genotypes. In *sut4* catkins, sucrose levels increased steadily during development, whereas in WT they remained stable (Fig. 5d). To assess whether the changes in *sut4* reflected increased vacuolar sequestration, we next examined sucrose-derived metabolites that originate in the cytosol.

Hexose levels increased in both genotypes during catkin expansion (Fig. 5d), but this is not a reliable indicator of cytosolic sucrose as hexoses can be derived from starch or sucrose pools in the vacuole, plastid, or cytosol. In contrast, biosynthesis of raffinose occurs exclusively in the cytosol and directly utilizes sucrose (Schneider and Keller, 2009). In WT, both galactinol and raffinose peaked in unopened floral buds and declined sharply during catkin development despite stable sucrose levels (Fig. 5d), a pattern consistent with high raffinose abundance in dormant poplar buds (Watanabe *et al*., 2018). However, this trend was absent in *sut4* trees. Raffinose levels were significantly lower in stage I *sut4* buds compared to WT, even though sucrose levels were similar. As catkins elongated and sucrose accumulated in *sut4*, raffinose, and to a lesser extent galactinol, also increased (Fig. 5d).

We also compared several Krebs cycle intermediates across all three stages of catkin development. Malic acid, the most abundant intermediate, increased throughout catkin development and did not differ between genotypes (Fig. 5d). In contrast, citric acid levels in *sut4* buds were less than half those of WT at stage I, partially recovered at stage II, and became comparable by stage III (Fig. 5d). This shift is notable given citric acid’s central role in fatty acid and acyl metabolism, and its connection to gluconeogenesis via the glyoxylate shunt under carbohydrate-limited conditions (Eastmond and Graham, 2001, Smith, 2002). Succinic acid, a product of citric acid metabolism during glyoxylate shunt activity, was significantly elevated in *sut4* catkins throughout development (Fig. 5d). These findings suggest that *SUT4*-KO affected metabolic pathways beyond carbohydrate metabolism.

### Transcriptional profiling revealed elevated stress responses in sut4 catkins

To investigate whether *sut4* infertility was due to insufficient protection during ovule development or to an inherent constitutive stress response, we performed RNA-Seq analysis on stage III elongating catkins. We identified 821 significantly up-regulated and 660 down-regulated genes in *sut4* catkins compared to controls (Dataset S2). Gene Ontology (GO) categories such as protein oligomerization, protein refolding, responses to heat, hypoxia, and hydrogen peroxide were significantly over-represented among the up-regulated genes (Fig. 6a). Interestingly, none of these GO categories were enriched among differentially expressed genes in vegetative tissues (e.g., leaves, xylem, bark, root tips) of *SUT4*-KD plants under benign greenhouse conditions (Xue *et al*., 2016). Specifically, over 22% of the ∼360 717 genes annotated to encode various heat shock proteins (HSPs) (Yer *et al*., 2018) were significantly up-regulated in *sut4* catkins. Also up-regulated were genes encoding several heat shock factors (HSFs) and late embryogenesis abundant (LEA) proteins. All three *galactinol synthase* genes detected in elongating catkins were sharply induced in *sut4* catkins, including the *Populus alba* × *grandidentata* ortholog previously shown to exhibit cool season-biased expression (Unda *et al*., 2012), but expression of *raffinose synthase* genes remained largely unchanged.

**FIGURE 6:**
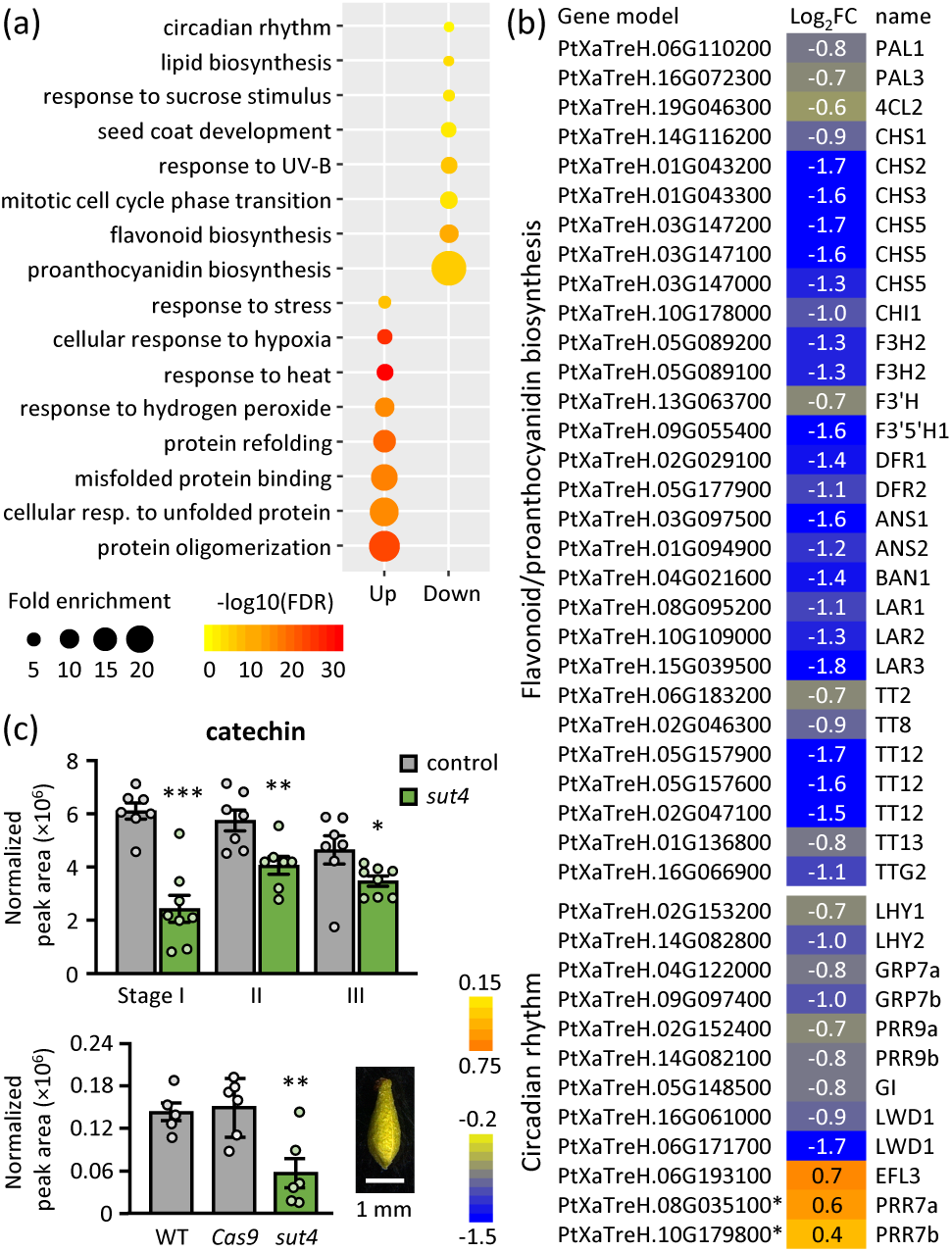
Gene expression responses in stage III elongating catkins. (a) GO enrichment of differentially up- or down-regulated genes in the elongating catkins of the *sut4* mutants relative to control. Representative GO terms with fold-enrichment >2 and gene number >5 are shown. (b) Heatmaps showing expression responses of genes associated with flavonoid and proanthocyanidin biosynthesis and regulation and circadian rhythm. Log_2_FC, log_2_-transformed fold-change (*sut4*/control) of *n* = 7 (pooled from 4 WT and 3 *Cas9*) and 5 (*sut4*) biological replicates. ANR, anthocyanidin reductase; ANS, anthocyanidin synthase; CHS, chalcone synthases; CHI, chalcone isomerase; DFR, dihydroflavonol reductase; EFL, early flowering; F3H, flavanone 3-hydroxylase; F3’H, flavonoid 3′-hydroxylase; F3’5’H, flavonoid 3′,5′-hydroxylase; GI, gigantea; GRP, glycine-rich RNA-binding protein; LAR, leucoanthocyanidin reductase; LHY, late elongated hypocotyl; LWD, light-regulated WD repeat-containing protein; TT, transparent testa; TTG, transparent testa glabra; PRR, pseudo-response regulator. (c) Catechin abundance during catkin development (top) or in pollinated carpellate flowers, with a representative image shown on the right (bottom). Data are mean ± standard error of *n* = 5-6 biological replicates. Significance was determined by Student’s *t*-test. *, *P* ≤ 0.01; ∗∗, *P* ≤ 0.01; ∗∗∗, *P* ≤ 0.001.

The smaller group of down-regulated genes were enriched for GO terms associated with flavonoid—particularly proanthocyanidin—biosynthesis, lipid biosynthesis, responses to UV-B and sucrose stimulus, mitotic cell cycle phase transition, seed coat development, and circadian rhythm (Fig. 6a). Nearly all flavonoid pathway genes (Tsai *et al*., 2006) and several upstream regulators detected in catkins were significantly down-regulated in *sut4* catkins (Fig. 6b). Also down-regulated were genes encoding orthologs of *Arabidopsis* TRANSPARENT TESTA12 (TT12), a multidrug and toxin extrusion (MATE)-type proton antiporter, and TT13 (also known as AHA10), a P-type H^+^-ATPase proton pump that energizes TT12-mediated proanthocyanidin transport into the vacuole (Debeaujon *et al*., 2001, Marinova *et al*., 2007). In *Arabidopsis*, both *TT12* and *TT13* are specifically expressed in ovules and developing seeds, and *tt12* and *tt13* mutants show defects in vacuole development and proanthocyanidin deposition in seed coat endothelial cells, resulting in reduced seed coat pigmentation and decreased dormancy (Debeaujon *et al*., 2001, Appelhagen *et al*., 2015). To examine whether *sut4* catkins exhibit reductions in proanthocyanidin abundance, we compared levels of the proanthocyanidin precursor catechin between genotypes. Consistent with the RNA-Seq data, catechin levels were significantly reduced during catkin development, as well as in individual carpellate flowers after pollination in *sut4* compared to controls (Fig. 6c).

Several genes encoding core components of the circadian clock system were significantly down-regulated in *sut4* catkins. These include *LHY1, LHY2* (late elongated hypocotyl), *GI* (gigantea), *GRP7a*, *GRP7b* (glycine-rich RNA-binding protein 7, also called CCR2 for cold circadian rhythm RNA-binding 2), *PRR9a* and *PRR9b* (pseudo-response regulator 9) (Edwards *et al*., 2018), whereas *PRR7a* and *PRR7b*, which act antagonistically with LHY (Nakamichi *et al*., 2007), were up-regulated (Fig. 6b). Many circadian clock genes are sensitive not only to light and temperature cues, but also to sucrose and abiotic stress signaling, influencing developmental, physiological, and metabolic functions (Haydon *et al*., 2017, Oravec and Greenham, 2022, Liang *et al*., 2024). Collectively, the transcriptomic data from catkins implicated elevated basal stress and circadian dysregulation as consequences of altered sucrose compartmentalization due to *SUT4*-KO.

## Discussion

### Knockout of SUT4 but not of SUT5/SUT6 accelerated autumn leaf senescence and delayed spring bud break and stem growth resumption

Seasonal alternation between carbohydrate storage during photoautotrophic growth and remobilization during heterotrophic subsistence is a hallmark of the perennial lifestyle (Regier *et al*., 2010, Tixier *et al*., 2019). Using CRISPR-induced frameshift mutants, we investigated the role of winter-expressed *SUT* plasticity during seasonal transitions. We found that CRISPR-KO of *SUT4*, which encodes a tonoplast-localized transporter for intracellular trafficking, but not *SUT5/SUT6*, which encode plasma membrane-localized transporter for export from the cell (Peng *et al*., 2014), had detrimental effects on leaf phenology and stem diameter growth in the field. The absence of phenotypes in *sut56* trees may reflect compensatory activity by SWEET15b, a plasma membrane-localized sucrose transporter with strong winter expression in poplar xylem (Ko *et al*., 2011, Zhang *et al*., 2021). In contrast, compensation for the loss of SUT4 appears less likely, as none of the SWEETs known to be expressed in poplar xylem encode tonoplast-localized sucrose transporters (Eom *et al*., 2015, Zhang *et al*., 2021).

*sut4* trees consistently exhibited premature leaf senescence, delayed spring budbreak, and delayed diameter growth relative to control and *sut56* trees (Fig. 1), effectively shortening their growing season. Leaf senescence typically spans 3-5 weeks (Fig. 1c), beginning with a sharp decline in chlorophyll and photosynthesis, followed by rapid cellular breakdown and nutrient resorption (Keskitalo *et al*., 2005). While *SUT5*/*SUT6* are weakly expressed in source leaves, *SUT4* is well expressed (Payyavula *et al*., 2011), which may explain the lack of seasonal phenotypes in *sut56* trees. A premature decline in photosynthetic capacity in *sut4* trees would be expected to lead to intracellular sucrose and turgor imbalances, further exacerbating the already elevated vacuolar sucrose sequestration. Although *SUT4*-KD or KO has been shown to alter leaf hydrostatics and pressure-volume characteristics (Harding *et al*., 2020), it does not affect leaf morphology or function under favorable growth conditions (Frost *et al*., 2012). Whether these physiological changes influence the timing or progression of autumn senescence remains unclear.

### Low raffinose may reflect metabolic or osmotic perturbations due to loss of SUT4 plasticity during winter

The early onset of senescence and premature leaf drop in *sut4* trees may have impaired the buildup of bud reserves and contributed to delayed spring bud break. Although vegetative buds were not analyzed, flowering bud data suggest that soluble sugar levels were not limiting in spring (Fig. 5d). Similarly, while spring diameter growth was delayed in *sut4* trees, summer growth rates were comparable to controls across multiple years (Fig. 1e, cohort 2). Notably, winter stem sugar and starch reserves were similar or even higher in *sut4* mutants (Fig. 2) and previous work has shown comparable sucrose and starch consumption rates in *sut4* and WT plants under carbon-limited conditions imposed by coppicing (Tuma *et al*., 2024). These findings suggest that spring growth delays may stem from factors other than reserve size or depletion rate.

Winter stem raffinose levels were much lower in *sut4* trees (Fig. 3). Given the antioxidant and osmoprotective roles of raffinose (Kornberg and Krebs, 1957, Regier *et al*., 2010), its attenuated accumulation may have contributed to delayed spring bud break and growth resumption. Among the possible causes are limited cytosolic sucrose (Schneider and Keller, 2009) and altered stress signaling upstream of raffinose biosynthesis (Pluskota *et al*., 2015, Khan *et al*., 2021, Noronha *et al*., 2022). Vacuolar sucrose sequestration in *sut4* trees could constrain RFO synthesis as seen in drought-stressed *SUT4*-KD leaves (Frost *et al*., 2012). SUT4 has also been implicated in stress modulation, with *SUT4*-KD and KO plants exhibiting changes in leaf pressure-volume curves, relative water content, and storage water capacitance during drought (Harding *et al*., 2020, Harding *et al*., 2022). A signaling role has also been suggested for galactinol, a precursor of raffinose, where its hyper-accumulation led to stunted growth and altered xylem properties in poplar (Unda *et al*., 2017). Together, these findings support a role for SUT4 in modulating sugar allocation and stress signaling, with consequences for seasonal growth.

### Perturbed sucrose and raffinose trajectories were associated with elevated stress during sut4 catkin development

Galactinol and raffinose are known to accumulate in dormant buds and developing seeds to confer desiccation and cold tolerance (Cox and Stushnoff, 2001, Peterbauer and Richter, 2001, Falavigna *et al*., 2018). Meiosis during gamete development is particularly stress-sensitive (Thakur *et al*., 2010) and raffinose has been shown to enhance floral bud hardiness (Cox and Stushnoff, 2001, Falavigna *et al*., 2018). During *sut4* catkin development, raffinose levels were low at stage I— when female meiosis is expected to have begun (Wang *et al*., 2012)—while WT buds peaked at this stage with multifold higher raffinose accumulation (fig. 5d). Since raffinose synthesis is modulated by cold and desiccation stress signaling (Peterbauer and Richter, 2001, Yan *et al*., 2022), the steady decline of galactinol and raffinose levels after stage I in WT is consistent with reduced stress. In contrast, *sut4* catkins showed a sustained increase, possibly reflecting delayed or abnormal metabolic activity under cytosolic sucrose limitation. Supporting this idea, *sut4* catkins had elevated succinic acid and reduced citric acid abundance during emergence and early elongation (Fig. 5d). One interpretation of this metabolic shift is increased gluconeogenesis to fuel dormancy release and reproductive development under sucrose-limited conditions (Eastmond *et al*., 2015, Watanabe *et al*., 2018, Fadón *et al*., 2020, Walker *et al*., 2021). Sustained raffinose levels may also indicate persistent stress in *sut4* catkins due to altered sucrose partitioning or stress signaling as discussed above. GO term enrichment for responses to heat, hypoxia, and hydrogen peroxide, along with protein oligomerization and refolding among genes upregulated in *sut4* catkins is consistent with heightened cellular stress (Fig. 6a).

High carbohydrate levels usually promote flavonoid biosynthesis, especially at low-temperature conditions (Araguirang and Richter, 2022). The widespread suppression of flavonoid biosynthesis pathway gene expression in *sut4* catkins is therefore counterintuitive given the elevated sucrose levels observed (Fig. 5d). This may suggest broader disruptions in cellular homeostasis. Impaired SUT4 symport activity for vacuolar sucrose efflux may have altered tonoplast membrane potential and affected other proton-dependent tonoplast transporters (Martinoia, 2017). Supporting this idea, the single-copy *AHA10/TT13* gene required for vacuolar proanthocyanidin transport (Appelhagen *et al*., 2015) was downregulated in *sut4* catkins. The coordinated down-regulation of *TT12*, *TT13*, and their upstream regulator *TTG2* which encodes a WRKY transcription factor (Gonzalez et al., 2016), alongside reduced catechin accumulation, points to cascading effects of SUT4-KO on vacuolar biogenesis and seed development. Concurrent up-regulation of heat shock proteins associated with protein oligomerization and refolding responses further supports a signaling role for SUT4 during catkin development.

### Loss of SUT4 function disrupted circadian clock gene expression in catkins

Flavonoid biosynthesis is positively gated by the circadian morning complex anchored by LHY (Harmer *et al*., 2000), and the circadian clock itself is modulated by sucrose, flavonoids, and various hormonal signals (Dalchau *et al*., 2011, Haydon *et al*., 2013, Haydon *et al*., 2017, Hildreth *et al*., 2022). In *sut4* catkins, morning- or daytime-expressed clock genes (*PtaLHYs*, *PtaGI*, *PtaPRR9s*) were downregulated, while evening- or nighttime-expressed genes (*PtaEFL3* and *PtaPRR7s*) were upregulated (Fig. 6b) (Takata *et al*., 2009, Ding *et al*., 2018, Toda *et al*., 2019). This pattern mirrors sucrose-modulated inverse regulation of *AtLHY* and *AtPRR7* in *Arabidopsis*, though *AtPRR9* was unresponsive (Haydon *et al*., 2013), unlike *PtaPRR9s*. Differential responses among orthologous PRRs are not surprising, given their evolutionarily conserved roles in clock regulation and documented species-specific expression divergence, which may reflect adaptive tuning to distinct environmental—particularly perennial—conditions. (Toda *et al*., 2019). It is worth noting that clock gene expression was unchanged in *SUT4*-KD plants under well-watered summer greenhouse conditions (Xue *et al*., 2016). This suggests that SUT4 plays a specific role in cool-season sugar sensing, especially in reproductive tissues where *SUT4* is highly expressed. Consistent with *sut4* phenotypes, altered phenology, including premature leaf senescence and bud set, delayed bud break, reduced growth and compromised freeze tolerance have been reported in transgenic poplar with downregulated expression of *LHY1*, *LHY2,* and *GI* (Ibáñez *et al*., 2010, Ding *et al*., 2018, Fataftah *et al*., 2021).

### SUT4 integrates sugar allocation, stress response, and circadian regulation

*SUT4* expression is responsive to abiotic stresses, including drought and heat, across multiple species (Xu *et al*., 2017), and its induction under cool-season conditions has been recently reported in poplar (Tuma *et al*., 2024). Mutation or downregulation of *SUT4* has been shown to affect biomass partitioning, flowering, and seed yield in rice (*Oryza sativa*), potato (*Solanum tuberosum*), and tomato (*Solanum lycopersicum*), with circadian gene expression changes observed in potato (Chincinska *et al*., 2008, Eom *et al*., 2011, Chincinska *et al*., 2013, Liang *et al*., 2023). The present work provides evidence supporting a role for SUT4 in regulating cytosolic sucrose availability during carbohydrate reserve depletion in winter stem and spring catkins of poplar. Loss of this capacity appears to have repercussions for both vegetative and reproductive growth, protective metabolism, and circadian clock function, highlighting the potential of SUT4 to coordinate sugar signaling, stress responses, circadian regulation, and seasonal development in poplar.

## Materials and Methods

### Production of *SUT-*KO lines

The generation of *sut4* mutants and Cas9-only vector controls in the *Populus tremula* × *alba* INRA 717-1B4 background were previously reported (Zhou *et al*., 2015, Harding *et al*., 2022). Two gRNAs were designed to target *PtaSUT5* and *PtaSUT6* individually to generate single mutants, and a third gRNA was designed to target the highly similar genome duplicates at a consensus sequence to generate double mutants. Target sites were cross-checked against the *Populus* VariantBD (Zhou *et al*., 2025) to ensure the absence of sequence polymorphisms. Oligo pairs (Table S1) were assembled into the p201N-Cas9 vector behind the *Medicago truncatula U6.6* promoter, following the approach of Bewg et al. (2022). *Agrobacterium-*mediated transformation and plant regeneration was performed as described (Bewg et al., 2022).

### Amplicon sequencing to determine mutation patterns

Leaves from independent events of tissue cultured plants were collected for genomic DNA extraction and amplicon sequencing library preparation (Bewg et al. 2022) using consensus primers (Table S1). Samples were then barcoded with Illumina amplicon indexing primers, pooled, and sequenced on an Illumina MiSeq at the Georgia Genomics and Bioinformatics Core at the University of Georgia to obtain PE150 reads. Sequencing reads were processed and indel patterns identified using the Analysis of Genome Editing by Sequencing (AGEseq) program (Xue & Tsai, 2015) with a mismatch allowance set at 1%, followed by manual curation. Transgenic events with biallelic (single KO) or tetra-allelic (double KO) edits were selected for further experiments (Dataset S1).

### Plant propagation and field establishment

Multiple independent lines for *sut4* (*n* = 8), *sut5* (*n* = 3), *sut6* (*n* = 2), and *sut56* (*n* = 3) mutants along with wild-type (WT) and Cas9 controls were propagated from single-internode stem cuttings (Frost *et al*., 2012). Cuttings were transferred to 1-gallon pots containing commercial soil mixture (Fafard 3B) supplemented with Osmocote (15-9-12 NPK, 3-month slow release) and placed in a walk-in growth chamber for ∼3 weeks (Tuma *et al*., 2024). Plants were moved into an adjacent greenhouse for 2 weeks before transfer to a lath house within a fenced nursery at the University of Georgia’s Whitehall Forest for an additional 2 weeks. For logistical reasons having to do with the timing of cutting establishment, *sut4* plants were slightly shorter than control plants at the time of out-planting. Plants were transferred to 20-gallon pots containing a mix of Jolly Gardener premium soil (75% aged pine bark, 25% peat moss) and Fafard 3B with additional Osmocote (15-9-12 NPK, 3-month slow release) and planted at the nursery in summer 2021. Genotypes were randomized throughout the site and spaced 1.5 meters apart from one another. The first planting occurred in early June 2021 for *sut4* mutants and controls (WT and *Cas9*), and the second planting occurred in mid-August 2021 for *sut4*, *sut5*, *sut6*, *sut56* and controls. Plants were watered using an overhead irrigation system for 2-4 hours daily from early April to late October. The field trial was conducted with approval from the Animal and Plant Health Inspection Service (APHIS) of the U.S. Department of Agriculture.

### Field growth and phenology monitoring

The diameter of the main stem ∼1m above the soil surface was measured monthly or bimonthly from August 2021 to April 2023, and after termination of most trees, monthly through August 2024 for the remaining trees. Plant height was measured from the base of the stem at soil level to shoot tip in April 2022 and 2023. Trees were monitored throughout the fall to compare onset and rate of leaf abscission between genotypes. Leaf color was compared between genotypes as an indicator of leaf senescence in October. In the spring, the appearance of bud flush on the dominant stem and flowering were recorded. Floral buds and catkin emergence were observed during spring of 2023 (late February) and 2024 (early March). The total number of trees that developed catkins was recorded prior to their removal from the field site per the APHIS regulation. Further information regarding the field trial monitoring, phenotyping, and sampling are shown in Fig. S1.

### Tissue sampling

Approximately 12–17 months after out planting, increment cores (10 mm in length and 2 mm in diameter) were collected from the main stem ∼1.3 meter above soil level using a micro-coring device (Costruzioni Meccaniche Carabin C, San Vito di Cadore, Italy) according to Rossi and colleagues (2006). Stems were approximately 2–6 cm in diameter at the time of sampling. Increment cores were collected from the same stems during active growth in August 2022 (summer) and during dormancy in January 2023 (winter). Xylem and bark were separated from the cores in the field, snap-frozen in liquid nitrogen before being transported back to laboratory facilities where they were stored at −80°C until sample processing occurred. Temperatures ranged from a night-time low of 22°C to a daytime high of 31°C (daily average 25°C) on the day of summer sampling, and from a low of 3°C to a high of 18°C (daily average 11°C) on the day of winter sampling (Fig. S1). Catkins were observed in late February 2023 after the second winter in the field. Three developmental stages of catkins from cohort 1 trees were collected on March 1, 2023 for transcript and metabolite profiling. All tissue sampling occurred under sunny conditions between 11AM – 2PM. Tissues were snap frozen, ground under liquid nitrogen with a mortar and pestle, and stored at −80°C.

### Controlled pollination

Branches containing flower buds were collected from WT and *sut4* trees and incubated in ¼MS macro salt solution at room temperature in a confined laboratory to force flower. After catkin flowers opened, wild *P. tremuloides* pollen sourced from Minnesota 2022 (with assistance of Kathy Haiby, Poplar Innovations Inc.) was brushed onto individual WT, *Cas9*, and *sut4* catkins, which were then covered with glassine bags. After 7-10 days, the capsules opened, and the cottony fuss was collected for seed observation under a dissecting microscope. The length of catkins, seed size and number, and ovule size were recorded. The absence of seeds from pollinated *sut4* flowers was first observed in 2023, and the controlled pollination experiments were repeated in 2024 and 2025 with reproducible results. Individual carpellate flowers collected 3-4 days after pollination in 2025 were flash-frozen in liquid nitrogen for catechin analysis by GC-MS as described below.

### Metabolite profiling

Tissues were lyophilized for 48 hours (FreezeZone 2.5, Labconco), ground through a 40-mesh sieve using a Wiley Mill (Thomas Scientific), and further ball-milled to a fine powder in a S1600 Mini G Bead-beater (Spex SamplePrep; Metuchen, New Jersey) for metabolite profiling by Gas Chromatography-Mass Spectrometry (GC-MS) as detailed previously (Harding *et al*., 2022, Tuma *et al*., 2024). Peak identities were assigned based on the NIST08, FiehnLib metabolomics library (Agilent) (Kind *et al*., 2009) and in-house spectral libraries from authentic standards (Xue *et al*., 2013, Chen *et al*., 2014, Harding *et al*., 2022). A standard mix of succinic acid, sucrose, glutamic acid, fructose, glucose, and ascorbic acid was loaded at the beginning and end of each sample sequence to monitor derivatization and instrument performance. Peak intensities were normalized by the internal standard and reported on a sample dry weight basis (normalized peak area). Sucrose and hexose abundance (sum of fructose and glucose) was reported as µg/mg dry weight based on calibration curves.

### Starch

Starch content was determined by hydrolyzing starch to glucose using enzymatic digestion with α-amylase and amyloglucosidase (Chow and Landhausser, 2004) using 10 mg of lyophilized, Wiley- and ball-milled tissue powder as previously described (Tuma *et al*., 2024). Starch abundance was reported as µg/mg dry weight based on a glucose calibration curve.

### Wood chemical composition and physical property analyses

Fifteen-cm long pieces of the main stem were destructively harvested following bud flush in March 2023. For wood chemistry analysis, stem segments were debarked, dried at 55°C for 24 hours, and ground to pass through a 40-mesh screen using a Wiley mill. Samples were then run through 99 cycles of hot ethanol extraction using a Soxhlet extraction unit (Buchi) and air dried. Total lignin content and the ratio of lignin monomers in the stems (syringyl/guaiacyl, S/G ratio) were determined by pyrolysis-molecular beam mass spectrometry according to Harman-Ware *et al*. (2022). Total lignin content was also determined by the Klason method as described (Swamy *et al*., 2015). Three ∼7.5 mm longitudinal wood sections were obtained from the main stem approximately ∼1 m above soil level for analysis of specific gravity, acoustic velocity, and modulus of elasticity as described (Vieilledent *et al*., 2018, Tsai *et al*., 2020, Dahlen *et al*., 2023).

### RNA-seq analysis

Snap-frozen catkin tissues from cohort 1 trees in 2023 were ground to a fine powder under liquid nitrogen and an aliquot was used for RNA extraction using the Direct-zol RNA Miniprep kit (Zymo) with PureLink Plant RNA Reagent (Life Technologies) as described (Harding *et al*., 2018). Illumina low input RNA (with PolyA selection) library preparation and NovaSeq sequencing (NovaSeq XP V1.5 reagent kits, 10B flowcell, 2 ×151 run length) were performed at the Joint Genome Institute (proposal: 10.46936/10.25585/6000884). RNA-seq reads were aligned to the *Populus tremula* haplotype of the 717 reference genome (Zhou *et al*., 2023) using STAR-2.7.10 (Dobin and Gingeras, 2015) with the following parameters: --alignMatesGapMax 20000, --alignIntronMax 10000, --outFilterScoreMinOverLread 0.1, and -- outFilterMatchNminOverLread 0.1. Expression levels were estimated using featureCounts from the Subread package v2.0.6 with the options -Q 2, -s 0, -T 12, -p, and --countReadPairs -C (Liao *et al*., 2014) and reported as FPKM (fragments per kilobase of transcript per million mapped reads). Differential gene expression analysis were performed using DESeq2 v1.34.0 (Love *et al*., 2014). Thresholds for significant differential expression were based on minimum FPKM ≥5 in all samples of at least one group, *Q* ≤0.005, and fold change ≥1.5. Gene ontology (GO) enrichment analysis of significantly up- or down-regulated genes was performed using ShinyGO v0.77 (Ge *et al*., 2020) available at http://aspendb.uga.edu/ShinyGO/. Expression responses of selected genes were visualized using the HeatMapperPlus tool (http://bar.utoronto.ca/ntools/cgi-bin/ntools_heatmapper_plus.cgi).

## Statistics

Repeated measures analysis of variance (ANOVA) was used to determine significant differences in growth over time, genotypic effects, and their interactions using *R* v4.1.0 (R Core Team, 2016) in Rstudio v.2022.12.353). All other reported genotypic contracts were assessed with Student’s *t*-test (α = 0.05).

## Data availability

The RNA-seq data from this study have been deposited to the National Center for Biotechnology Information Sequence Read Archive under accession numbers SRP540970 to SRP540981.

## Acknowledgements

The authors thank Gilles Pilate of INRA, France for providing poplar clone INRA 717-1B4, Batbayar Nyamdari, Aitebiremen Gift Omokhua-Uyi, and Nuoendagula for GC-MS assistance, Maria Ortega for guidance on pollination experiments, Hansini Fernando for assistance with sample processing, the Georgia Genomics and Bioinformatics Core at the University of Georgia (RRID:SCR_010994) for Illumina sequencing assistance, the Georgia Advanced Computing Resource Center for computational resources, and Kathy Haiby of Poplar Innovations Inc. for aspen pollen collection.

## Author Contributions

TTT, SAH, and CJT conceived the study and designed all experiments. TTT performed all experiments and analyzed the data, with assistance from HAM, HP, and WPB for transgenic plant production and amplicon sequencing, SAH for metabolite data analyses, BL for RNA extraction, AL, KWB, and RZ for RNA-seq analysis, DLWW for diameter and flower monitoring, MD and RZ for controlled pollination, JD for wood property analysis, and SS and AEHW for wood chemistry analysis. TTT prepared the manuscript which was revised by CJT with contributions from SAH.

## Funding

This work was supported in part by a National Science Foundation Graduate Research Fellowship (DGE 1842396) to T.T.T., a Department of Energy, Office of Science, Biological and Environmental Research Program Award (DESC0023166) to C.-J.T., the Center for Bioenergy Innovation, Department of Energy, Office of Science, Biological and Environmental Research Program (Award Number ERKP886) to C-J.T and A.E.H.-W. The work (proposal: 10.46936/10.25585/6000884) conducted by the U.S. Department of Energy Joint Genome Institute https://ror.org/04xm1d337), a DOE Office of Science User Facility, is supported by the Office of Science of the U.S. Department of Energy operated under Contract No. DE-AC02-05CH11231. This work was authored in part by the Alliance for Sustainable Energy, LLC, the manager and operator of the National Renewable Energy Laboratory for the US Department of Energy (DOE) under contract no. DE-AC36-08GO28308. The views expressed in the article do not necessarily represent the views of the DOE or the US government. The US government retains and the publisher, by accepting the article for publication, acknowledges that the US government retains a nonexclusive, paid-up, irrevocable, worldwide license to publish or reproduce the published form of this work, or allow others to do so, for US government purposes.

## Conflict of interest disclosure

none

## Supporting Information

Table S1. Primers

Figure S1. Field trial timeline and temperature profiles

Dataset S1. Mutation patterns of *SUT5*, *SUT6*, and *SUT56* edited transgenic lines.

Dataset S2. Differentially expressed genes

## Notes

### Competing Interest Statement

The authors have declared no competing interest.

### Summary of Updates

Added new data and revised the introduction and discussion sections significantly

